# An unconventional myosin, *myosin 1d* regulates Kupffer’s vesicle morphogenesis and laterality

**DOI:** 10.1101/268789

**Authors:** Manush Saydmohammed, Hisato Yagi, Michael Calderon, Madeline J. Clark, Timothy Feinstein, Ming Sun, Donna B. Stolz, Simon C. Watkins, Jeffrey D. Amack, Cecilia W. Lo, Michael Tsang

## INTRODUCTION

Establishing left-right (LR) asymmetry is a fundamental process essential for arrangement of visceral organs during development. In vertebrates, motile cilia driven fluid flow in the left-right organizer (LRO) is essential for initiating symmetry breaking event^1–3^. Without a definite LRO structure in invertebrates, LR asymmetry is initiated at a cellular level by actin-myosin driven chirality^4, 5^. In *Drosophila*, myosin1D drives tissue-specific chirality in hind-gut looping^6, 7^. Here, we show that *myosin 1d* (*myo1d*) is essential for establishing LR asymmetry in zebrafish. Using super-resolution microscopy, we show that the zebrafish LRO, Kupffer’s vesicle (KV), fails to form proper lumen size in the absence of *myo1d*. This process requires directed vacuolar trafficking in KV epithelial cells. Interestingly, the vacuole transporting function of zebrafish Myo1d can be substituted by myosin1C derived from an ancient eukaryote, *Acanthamoeba castellanii*, where it regulates the transport of contractile vacuoles. Our findings reveal an evolutionarily conserved role for an unconventional myosin in vacuole trafficking, lumen formation and determining laterality.

## RESULTS

Several animal models have been used to investigate different theories for establishment of LR asymmetry^8^. This includes intracellular chirality^9^, voltage gradient flow^10^, chromatid segregation^11^ and motile cilia^12^. Cilia-driven fluid flow has been reported to be essential for LR asymmetry in *Mus musculus* (mouse)^1^, *Xenopus laevis* (African Clawed Frog)^3^, *Danio rerio* (zebrafish)^2^ and *Oryctolagus cuniculus* (rabbit)^13^. However, a role for cilia is not universally conserved across vertebrates. In chick embryos, symmetry breaking is regulated by asymmetric cell migration around Hensen’s node and does not involve cilia based flow^14^. In addition, motile cilia are not associated with LR asymmetries that develop in invertebrates, such as the asymmetric looping of the gut and gonad in *Drosophila*^6, 7^. Therefore, probing common cellular events in different model systems may reveal insights into the early mechanisms of breaking symmetry that have been conserved during evolution.

Prior studies involving *Drosophila* myosin1D revealed a role for actin based molecular motors in establishing LR asymmetry in invertebrates^6, 7^. We wondered whether *myo1d* has an evolutionary conserved role in specifying laterality in vertebrates. Zebrafish *myo1d* is expressed early and ubiquitously, including within tailbud region where the ciliated KV forms (white arrow, Supplementary Fig 1a). We generated *myo1d* mutants using goldy TALENs (Supplementary Fig. 1b)^15,16,17^ to reveal its role in LR patterning. Three mutant alleles of *myo1d*, herein named *pt31a, b* and *c* (Supplementary Fig. 1c) were predicted to cause frame shift and amino acid truncation (Supplementary Fig. 1d). However, zygotic homozygous mutants (*myo1d*^*p31a/pt31a*^) survived to adulthood in a Mendelian ratio suggesting that *myo1d* maternal expression was sufficient for embryonic development. We generated *myo1d*^*pt31a/pt31a*^ maternal-zygotic (MZ) mutants and found that the KV had either a small or dysmorphic lumen at 8 somite stage (S) (Fig. 1 ai-iii). Consistently, injection of *myo1d* translation blocking antisense morpholinos also affected KV size (Supplementary Fig. 2a, b & Supplementary Movie 1 & 2). Next, we stained for tight junction protein, ZO-1 to assess KV morphology in *myo1d* MZ mutants. ZO-1 staining showed a smaller or dysmorphic KV lumen compared to controls (Fig. 1 aiv-avi), indicating *myo1d* is essential for generating a spherical KV shape and forming a lumen. We crossed *myo1d*^*pt31a*^ mutants into a transgenic line that labels membranes of KV cells, *Tg*(*dusp6:EGFP*)^*pt21*^, and confirmed that *myo1d* MZ mutants have smaller or dysmorphic lumen (Fig. 1a vii-ix, & 1b). Next, we confirmed that Myo1D expression was detected in KV epithelial cell borders (Supplementary Fig. 3a), whereas it was absent in *myo1d* MZ KVs (Supplementary Fig 3b), suggesting that the *pt31a* allele is a null mutant. Also, a similar lumen defect was observed in the otic vesicle (OV) (Fig. 1c-e), suggesting that *myo1d* is essential for the formation of different fluid filled structures during development. A report has shown that a minimum threshold lumen size is necessary for establishing robust LR asymmetry in zebrafish^18^. Consistently, we found increased frequency of laterality defects in *myo1d* MZ mutants (Fig. 1f & g) and in *myo1d* morphants over wildtype (Supplementary Table 1). These experiments revealed a role for *myo1d* in establishing proper KV shape, lumen size and laterality in zebrafish.

**Figure 1.**
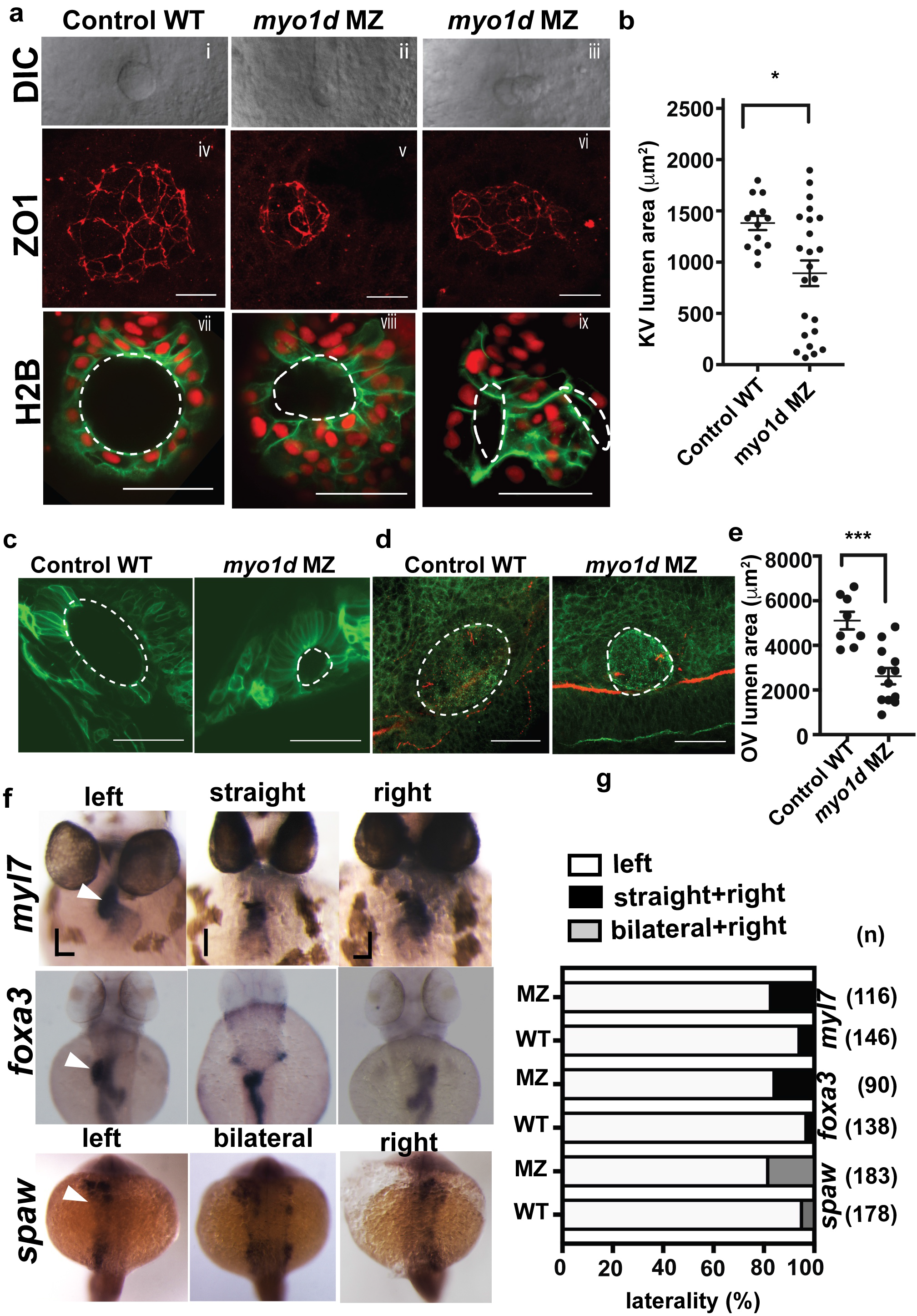
Myold is necessary for the formation of kupffer’s vesicle lumen and determining laterality. (**a**) *myo1d* mutants showed KV lumen formation defects. **i-iii**, DIC images. (**iv-vi**), ZO-1 tight junction staining. **vii-ix**, *Tg*(*dusp6:EGFP*)^*pt21*^ embryos injected with H2B mCherry mRNA showing KV cell and nuclei, respectively. *myo1d* MZ mutant KVs were either with smaller or dysmorphic lumen. Lumen area outlined with white dotted markings. (**b**) Graph showing KV lumen area in *myo1d* MZ mutants (n=23) compared to wildtype (n=13). (**c**) Otic vesicle lumen defects at 24hpf in *myo1d* MZ embryos was detected when compared to wildtype embryos. *Tg*(*dusp6:EGFP*)^*pt21*^ was used to delineate otic vesicle lumen membrane (dotted line). (**d**) aPKC (green) and acetylated tubulin (red) antibodies were used to show otic vesicle morphology. **e**, Representative graph showing otic vesicle (OV) lumen area of WT (n=8) and *myo1d* MZ mutants (n=12). (**f**) *myo1d* MZ mutation affected laterality of the heart (*myl7*), endoderm (*foxa3*) and lateral plate mesoderm (*spaw*). (*g*) Graph showing percentage of embryos with laterality defects in *myo1d* mutants (MZ) and wildtype (WT). n represents number of embryos scored. Two sample comparisons were derived from unpaired Students t-test and Mann Whitney test. * p<0.05, ***p<0.0001 represent a statistical difference. Data as mean ± SEM. (**avii-aix, c**) single plane Z-stack images showing the largest lumen area while, **d** are 3D projection images.

Since motile cilia in the KV are required for proper laterality, we determined if loss of *myo1d* affected ciliogenesis. Acetylated tubulin staining showed that cilia length was normal (Supplementary Fig. 4a & b) but motile cilia numbers per KV were less in *myo1d* MZ mutants (Supplementary Fig. 4c). As KV epithelial cells are monociliated^2^, we counted KV nuclei bordering the lumen and found fewer cells in *myo1d* MZ embryos (Supplementary Fig. 4d & e). Thus, the decreased cilia number was due to fewer KV cells. Together, these results indicated that *myo1d* loss has no effect on ciliogenesis.

At 6 hours post fertilization (hpf), KVs are formed from the dorsal forerunner cells (DFCs) near the organizer^2^. These cells proliferate and undergo a mesenchymal-to-epithelial transition to form a rosette like structure, where KV lumen forms^2^. To assess if the abnormal KV lumen phenotype in *myo1d* MZ mutants are a result of defects in DFC clustering, we analyzed *foxj1a* expression^19^. We did not observe differences in *foxj1a* expression in *myo1d* MZ and wildtype embryos (Supplementary Fig. 5). This indicated that DFC clustering and migration was not the cause of defective KV morphogenesis.

KV lumen formation is a rapid and dynamic process during somitogenesis that spans approximately 3-hour period (1S to 8S, 10-13 hpf) when a fluid filled spherical shaped organ is aligned to the base of the notochord (see Supplementary Movie 1)^2^. Previous studies described how a symmetric KV rosette undergoes extensive cellular remodeling to become an asymmetric KV so that anterior cells retain columnar shape whereas posterior cells attain squamous or cuboidal shape^20^. On depletion of Rock2b or pharmacological inhibition of non-muscle myosin II was found to affect anterior-posterior (AP) cell shape changes in the KV^21^, suggesting that actomyosin activity is important for generating AP asymmetry. Additional fluid filling mechanisms may also be contributing to this process.

Thus, we reasoned that *myo1d* could contribute to the AP asymmetry through a fluid filling mechanism. In *Tg*(*dusp6:EGFP*)^*pt21*^ embryos, we observed vacuole-like structures in KV cells that were designated as such by virtue of their size (Fig. 2ai, aii, Supplementary Movie 3 & 4), and were larger than endocytic or ER derived vesicles^22^. These vacuoles were noted in the KV suggesting these structures are the basis for expanding the KV lumen. To quantify this, we counted number of vacuoles in KV cells during lumen expansion. At 3S, vacuoles appeared near the plasma membrane of the every KV epithelial rosette cell (Fig. 2ai). By 8S, the number of vacuoles was significantly less in posterior cells compared to 3S (Fig. 2aii, & b). In *myo1d* MZ mutants and *myo1d* AUG morphants, we observed an increased number of vacuoles in posterior KV epithelial cells at 8S (Fig. 2 aiii-iv & c) and posterior KV cells remained columnar shape. Consistently, transmission electron microscopic images revealed an abundance of vacuole-like structures in the *myo1d* MZ KVs (Fig. 2d). Moreover, Myo1d was colocalized with vacuoles in KV cells (white arrow, Supplementary Fig. 3a), suggesting Myo1D is involved in trafficking of these vacuoles. Together, these experiments suggest that vacuole clearance contributes to AP cell shape changes in the KV.

**Figure 2.**
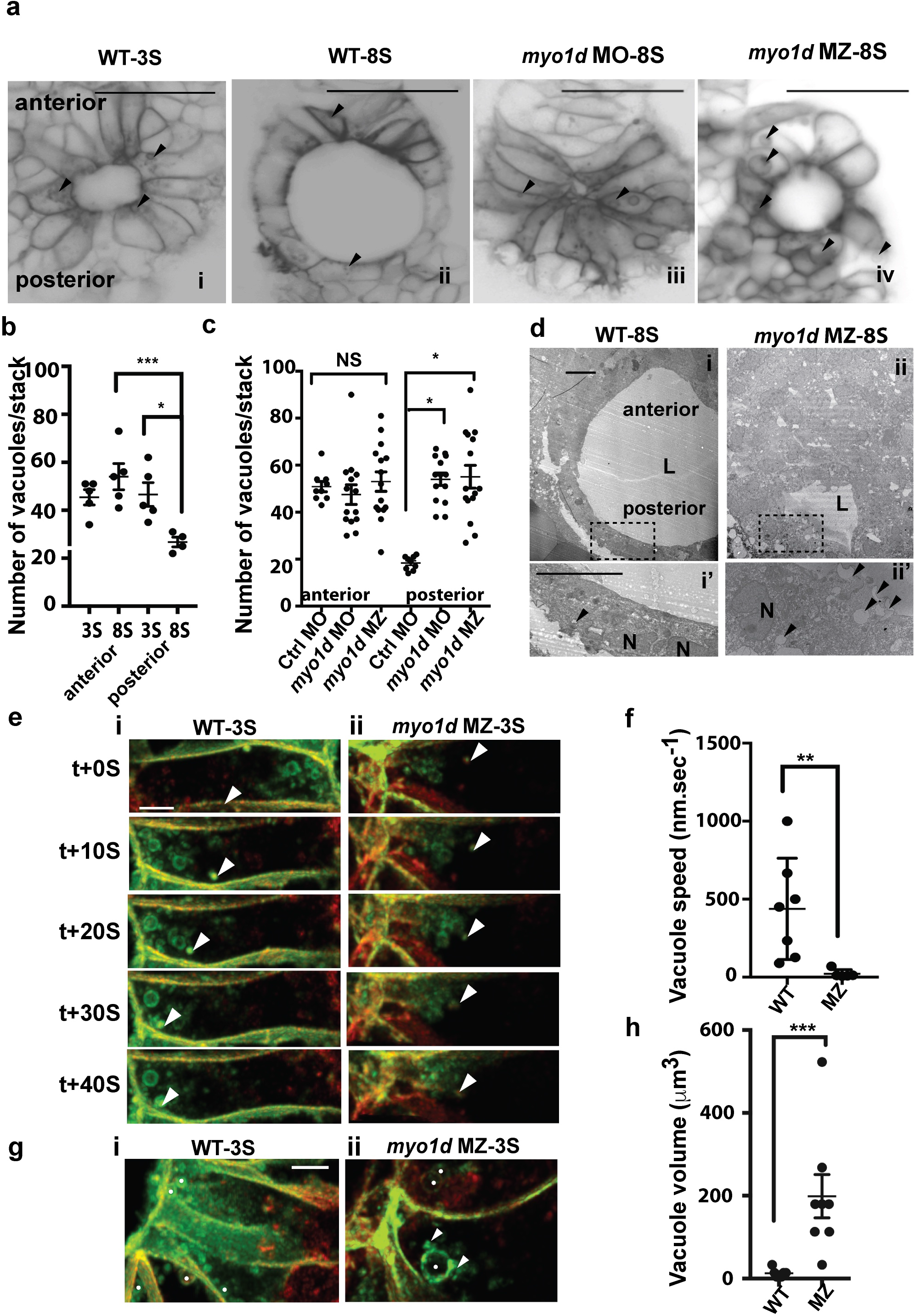
Myold mediates directed vacuolar movement in KV cells. (**a**) Predominance of vacuoles in KV epithelial cells (black arrowheads) from 3S (**ai**) and 8S (**aii**). At 8S, *myo1d* morphants (**aiii**) and *myo1d* MZ mutants (**aiv**) showed an abundance of large vacuoles in KV cells. (**b**) Graph showing vacuole counts in anterior and posterior region between 3S (n=5) and 8S (n=5) stage. (**c**) Graph showing vacuole counts in anterior vs posterior region in control MO (n=9) myo1d MO (n=14) and *myo1d* MZ (n=15) (**d**) TEM showing the presence of vacuole (black arrowhead) in posterior KV cells was less in wildtype (**di-i’**, n=2) than *myo1d* MZ (**dii-ii’**, n=2). L-lumen, N-nucleus. (**e**) 4D STED time-lapse images of a KV cell showing directed vacuolar migration towards apical surface (white arrowhead) (n=3). Vacuole movement were slower in the *myo1d* MZ mutants (**eii**, n=3) compared to wildtype (**ei**, n=3). (**f**) Average vacuole movement speed decreased in *myo1d* MZ mutants (nv=7) over wild type (nv=5). (**g**) KV epithelial cell showing small vacuole size in wildtype (**gi**) and compared to vacuole size in *myo1d* MZ (**gii**) mutants. White spots mark vacuoles. (**h**) Average vacuole volume increased in *myo1d* MZ (nv=7) when compared to wildtype (nv=8). Note small vacuoles (white arrowheads) fusing to large vacuole (white spots) in **gii**. Statistical comparisons in **b & c** by ANOVA and post hoc analysis with Turkey’s multiple range tests, whereas for **f & h** by unpaired Students t-test and Mann Whitney test. * p<0.05, =*p<0.01, ***p<0.005, represent a statistical difference. NS, not significant. Data as mean ± SEM. nv-number of vacuoles analyzed, n- number of KV analyzed. scale bars: (**a,b**)-50 μm, **e**-10 μm, (**f,g**)-5μm.

Next, we used super resolution imaging to precisely track vacuolar movement from 2-3S. The epithelial cells lining KV showed directed vacuolar trafficking towards the rosette apex (Fig. 2ei & Supplementary Movie 5). This vacuolar movement is dynamic such that smaller vacuoles can fuse together as they reach the KV apical membrane (Fig. 2ei, & Supplementary Movie 5). On the contrary, *myo1d* MZ embryos revealed a dispersed, largely cytoplasmic distribution of vacuoles and their movements were misdirected (Fig. 2eii). Moreover, the speed of vacuolar movement was decreased in *myo1d* MZ mutants (Fig. 2eii, f & Supplementary Movie 6). Continuous vacuole fusion events (white arrow Fig. 2g) resulted in larger vacuoles (compare vacuoles marked with white spots in Fig. 2gi vs gii, Supplementary Movie 5 & 6). Further, larger vacuoles were predominant in KV epithelial cells of *myo1d* MZ mutants (Fig. 2h, Supplementary Movie 7 & 8). Thus, *myo1d* is required to deliver fluid filled vacuoles to the KV lumen apex. These observations establish the role of *myo1d* in directed vacuolar movement as a mechanism for KV lumen expansion.

In ancient unicellular eukaryotes, myosin-I is involved in transporting water through contractile vacuoles (CV), which is essential for attaining amoeboid cell shape and regulating directed cell motility^23,24^. Loss of *Acanthamoeba castellanii* myosin-IC activity by inhibitory antibodies resulted in CV accumulation in the cell that ultimately leads to cell rupture^23^. We reasoned that a similar process may be occurring in the KV epithelial cells in *myo1d* MZ mutants. We found significantly higher number of fragmented nuclei in *myo1d* MZ embryos compared to wildtype at 1S (Fig. 3a, & b) and 6S stage (Fig. 3c & d), suggesting that accumulation of vacuoles may cause KV epithelial cells to burst that also account for the lower cell number (Supplementary Fig. 4d & e). We postulated that an *Amoeba* Myosin-1C could compensate for the loss of *myo1d* in zebrafish. Overexpression of *Acanthamoeba* myosin-IC expanded KV lumen in wildtype embryos (Fig. 3e, f), indicating that myosin-IC derived from *Amoeba* is functional in zebrafish KV cells. Moreover, injection of *Acanthamoeba* myosin-IC mRNA into *myo1d* MZ embryos rescued KV lumen and AP cell morphology defects (Fig. 3g & h) as efficiently as a phylogenetically closer version, *Myosin 1d* derived from rat (Supplementary Fig. 6a &b). Together, class I myosins play an evolutionary conserved role in intracellular vacuolar transport that regulates cell shape changes and drives lumen formation in the zebrafish KV.

**Figure 3.**
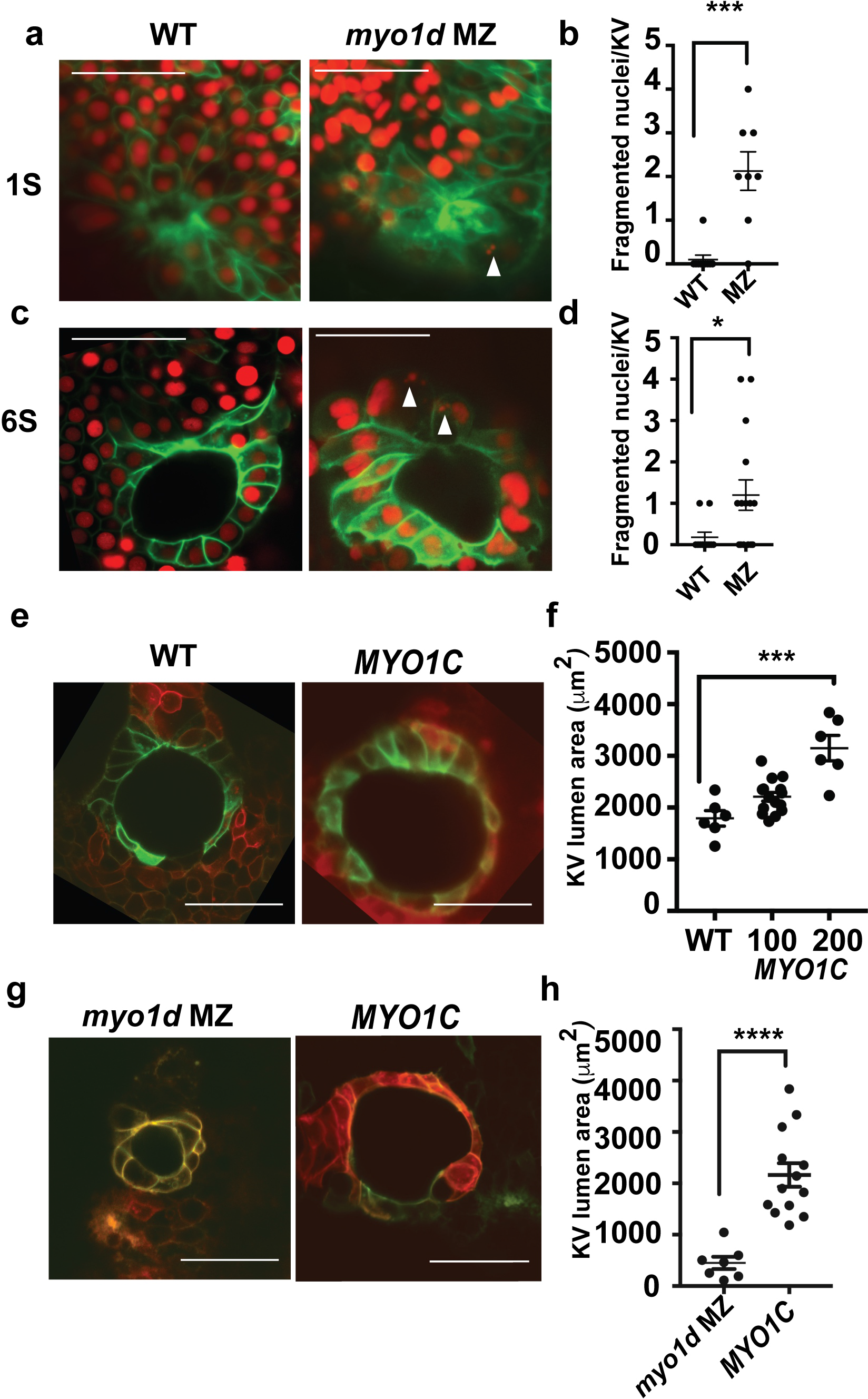
Substituting *myold* with amoeba myosin 1C rescues KV lumen defects. (**a**) Fragmentation of nuclei (white arrowheads) in wildtype or *myo1d* MZ KV at 1S. (**b**) Graph showing increased fragmented nuclei in *myo1d* MZ (n=8) mutants over wildtype (n=10). (**c**) Fragmentation of nuclei (white arrowheads) in wildtype and in *myo1d* MZ KV at 6S. (**d**) Graph showing fragmented nuclei in *myo1d* MZ (n=15) KVs and in wildtype controls (n=11). (**e**) Overexpression of *Acanthamoeba* myosin-IC was sufficient to expand lumen area in wildtype KV. (**f**) Graph showing KV lumen area after overexpression of *Acanthamoeba* myosin-IC (100pg; n=15, 200 pg; n=6) compared to wildtype KV (n=6). (**g**) Rescue of myo1d MZ mutant KV with *Acanthamoeba* myosin-IC. **h**, Graph showing KV lumen area in myo1d MZ mutants (n=7) and after *Acanthamoeba* myosin-IC mRNA injections (n=13). For **a-d**, *Tg*(*dusp6:EGFP*)^*pt21*^ embryos injected with H2B-mCherry mRNA. In **e & g**, 50 pg pCS2-CAAX mCherry as injection control. Two sample comparisons in **b, d & h** were by unpaired Students t-test and Mann Whitney test. For **f**, One-way ANOVA and post hoc analysis with Turkey’s multiple range tests. * p<0.05, ***p<0.001, ****p<0.0001 represent a statistical difference. Data as mean ± SEM. n number of KV analyzed. scale bar-50μm.

Cystic fibrosis transmembrane conductance regulator (CFTR), which is localized in KV apical membrane, was shown to regulate water transport into the epithelial lumen^25^. Consistently, zebrafish CFTR mutants exhibited impaired KV lumen formation^26^. Also, pharmacological treatment with Forskolin and IBMX (FIBMX) that activates CFTR expands KV lumen size^27^. We questioned whether CFTR can still function in *myo1d* MZ mutants. Quantification of CFTR localization to the KV apical surface was performed using the *TgBAC*(*cftr-GFP*) embryos^26^ (Supplementary Fig. 7). We observed normal CFTR localization in the KV apical membrane after Myo1d depletion (Fig. 4a, b). Interestingly, FIBMX treatment in *myo1d* MZ embryos showed expansion of KV lumen, implicating CFTR mediated lumen expansion was still functional in the absence of Myo1D (Fig. 4c-e). These results support a model where *myo1d* and CFTR are two independent mechanisms regulating the fluid filling process in the KV (Fig. 4f). Thus, proper KV lumen formation requires multiple modes of fluid filling mechanisms such that a threshold volume is achieved in a limited developmental timeframe. It appears that KV fluid is primarily derived from posterior epithelial cells that decrease cell volume concomitant with lumen expansion. It is remarkable that this process is akin to the water expulsion mechanism found in protozoans that traffic water and other fluid filled contractile vacuoles to the plasma membrane. Similar asymmetrical fluid loss and epithelial thinning process were observed during zebrafish otic vesicle development where actomyosin interaction provides the forces necessary for expansion of lumen^28^. Consistently, this remodeling process creates new luminal space and cause a net redistribution of fluid from epithelial cells to lumen, highlighting the role of intra epithelial fluids in lumen expansion^28^. We also found lumen formation defects in the otic vesicle of *myo1d* MZ mutants (Figure 1c-e), suggesting that *myo1d* mediated epithelial cell thinning process could be a common mechanism for lumen formation during development.

**Figure 4.**
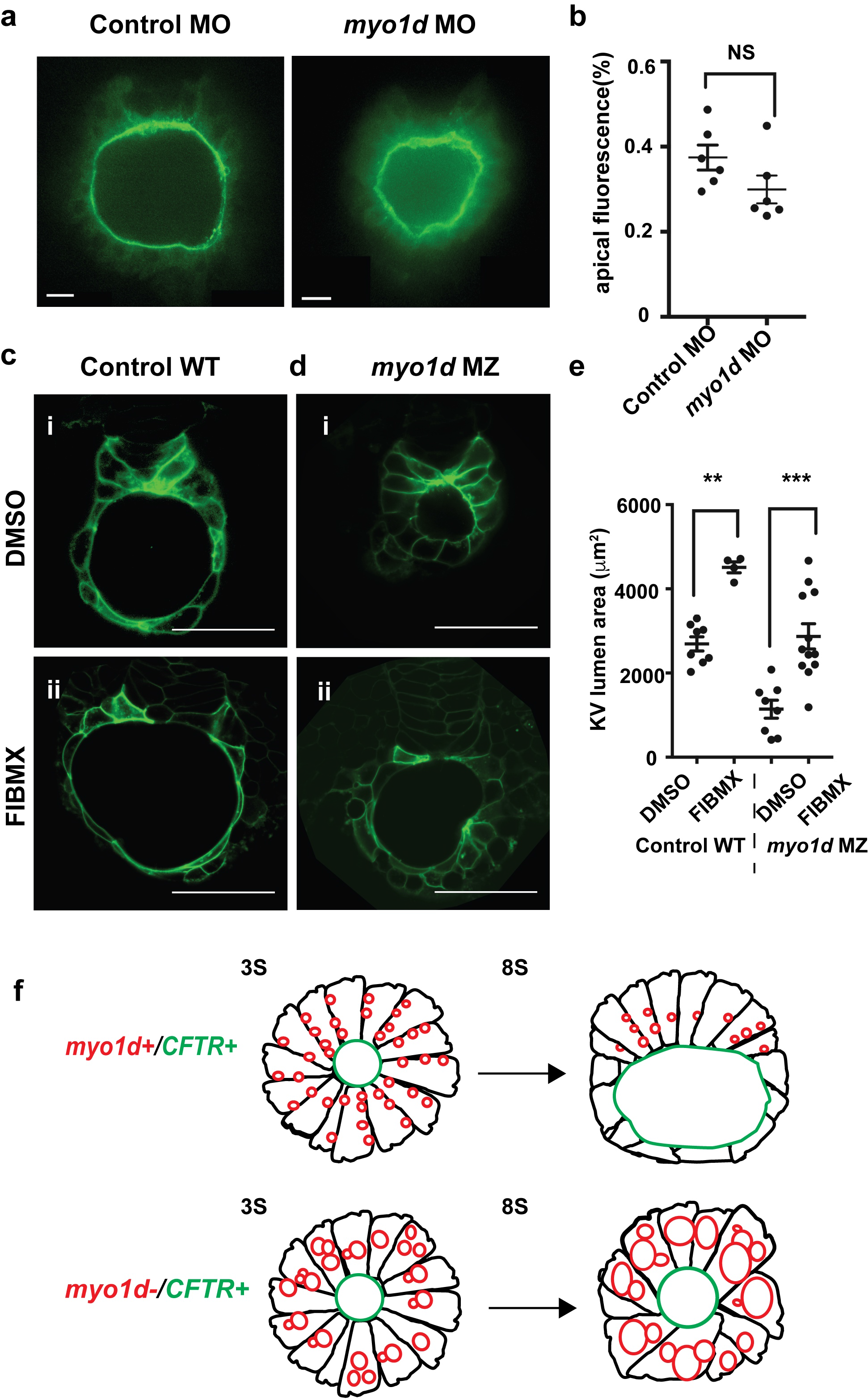
Loss of *myold* does not affect cAMP activation and CFTR activity. (**a**) Control MO or *myo1d* MO injected *TgBAC*(*cftr-GFP*) embryos did not affect CFTR apical GFP expression. (**b**) Graph showing CFTR apical fluorescence was not affected in *myo1d* MO injected(n=6) compared to wildtype KV (n=6). (**c**) FIBMX treatment (**cii**) increased lumen size as compared to DMSO (**ci**) in wildtype KV. (**d**) FIBMX treatment rescued lumen size in *myo1dMZ* (**di**) embryos when compared to DMSO (**dii**) treatment. (**e**) Graph showing FIBMX treatment in both wildtype and *myo1d* MZ embryos significantly expands lumen volume. wildtype DMSO (n=8), wildtype FIBMX (n=4), *myo1d* MZ DMSO(n=8), *myo1d* MZ FIBMX (n=12). (**f**) A proposed model depicting the role of Myo1d and CFTR in KV lumen expansion. When Myo1d and CFTR channel are active, proper AP cell morphology and threshold lumen expansion occurs by the 8S. In the absence of Myo1d, despite the presence of CFTR channel, KV lumen fails to expand. This is due to the failure of vacuole clearance from KV epithelial cells. For **c & d**, IBMX (10μM), and Forskolin (40μM) used from bud to 8S. (n=12). For **a, c & d**, KV imaged at 8S. For **b**, statistical comparisons using unpaired Students t-test and Mann Whitney test, whereas in **d**, One-way ANOVA and Turkey’s multiple range tests. Not significant-NS, **p<0.001, ***p<0.0001 represent a statistical difference. Data as mean ± SEM. n-number of KV analyzed. scale bars: **a**-15 μm, **c-d**-50 μm.

In *Drosophila*, tissue specific and temporal expression of myosin1D in the hindgut was sufficient to drive intrinsic chirality at the cellular level that generates a consistent gut and gonad looping pattern^4^. A mechanical model suggests that anchored myosin motors walking along actin drives the filaments to turn in leftward circles^29^. With no LRO in invertebrates, these circular forces drive specific organ looping morphogenesis^4, 5^. On the contrary, in zebrafish, generation of AP cell shape changes in the KV is the earliest asymmetric event. *myo1d* contributes to AP asymmetry in the zebrafish, prior to ciliogenesis and establishment of fluid flow^6, 7^. Together, we reveal a conserved vacuolar transport function of a myosin motor protein found in primitive eukaryotes that is critical in determining left-right asymmetry in a vertebrate.

## MATERIALS AND METHODS

### Zebrafish handling and maintenance

All the experiments using zebrafish was carried out with prior review and approval by the University of Pittsburgh Institutional Animal Care and Use Committee. Transgenic zebrafish lines used in work were: *Tg*(*dusp6:EGFP*)^*pt21*^ AB*, *myo1d* TALEN mutant lines generated for this work were labeled: *pt31a*, *pt31b* and *pt31c* alleles.

### Morpholino injections

Antisense Morpholino (MOs) were obtained from Gene-tools LLC. MOs were designed against translation initiation codon (5’-CCAAACTTTCGTGTTCTGC**CAT**AAT-3’) of zebrafish *myo1d*. MOs were injected into embryos at 2-cell stage as previously reported^30^. To conduct rescue experiments, full length rat MYO1D was PCR amplified and cloned into pCS2+ vector. *Acanthamoeba* sp myosin-1C^23^ was cloned into pCS2+ vector between Nco1 and Xba1 restriction site. Linearized pCS2+ vector was used to generate mRNA using SP6 mMessage mMachine (Ambion: AM1340) and purified using Roche mini quick spin RNA columns (Roche: 11814427001, USA.).

### Design of TALEN DNA binding domains

The software developed by the Bogdanove laboratory (https://boglab.plp.iastate.edu/node/add/talen) was used to find best DNA binding sites to target *myo1d* exon2 as described^15^. TALEN binding sites were 18 (Random variable repeats) RVDs (NI HD HD NH NG NH HD HD NI NG NH NI NI HD NI NG NG NG) on the left-TALEN and 19 RVDs (HD NG NH HD HD NG NG NG NH NG NI HD NG NH HD NG HD NI NI) on the right-TALEN with 15 bases of spacer.

### Construction of goldy TALENs and mutant generation

TALEN assembly of the RVD-containing repeats was conducted using the Golden Gate approach^31^. Once assembled, the RVDs were cloned into a destination vector, pT3TS-GoldyTALEN. After sequence confirming RVDs were in frame with destination vector, linearized plasmids were used to generate mRNAs using SP6 mMessage mMachine kit (Ambion: AM1340), and purified using Roche mini quick spin RNA columns (Roche: 11814427001, USA). Each mRNA TALEN pairs were co-injected into 1-cell stage embryos and raised to adults. Somatic and heritable TALEN induced mutations were evaluated by using forward (5’-TTGCTGCAGGTTTGAAAAGGGTCGTA-3’) and reverse (5’-CAGTCTACCTGAGATGACAATGCACG-3’) using primers and One Taq DNA polymerase (NEB: MO0480S). PCR was performed using following cycling conditions: initial denaturation 4 min 95 °C, 35 cycles (30S 95 °C, 30S 58 °C, 30S 68 °C) and final extension 5 minutes 68°C. A 226 bp PCR amplicon generated from 48hpf embryos were used to detect disruption of AatII restriction site in the *myo1d* target site by restriction fragment length polymorphisms (RFLPs). Founders harboring mutant alleles were outcrossed to generate F1 population and sequence verified. Positive F1 heterozygotes were in-crossed to generate homozygous zygotic (Z) mutants. Maternal zygotic (MZ) mutants are generated by in-crossing homozygous parents.

### Riboprobes preparation and in situ expression analysis

Total RNA was isolated from zebrafish embryos at 24hpf. Total RNA (1 μg) was reverse transcribed with Superscript II Reverse Transcriptase (Invitrogen) and amplified with the primers: 5’-AACGTTCCTCCTTGCCCTGTAATC-3’ (forward) and 5’-TCTATAATGTGACCGGAGTGAGCA-3’ (reverse). PCR products of *myo1d* mRNA and inserted into PCRII-TOPO vector (Thermo Fisher Scientific: K465001). Sequence confirmed clones were used to make *myo1d* riboprobes. Following riboprobes prepared using the cDNA constructs for *myl7* ^32^, *spaw* ^33^ *foxa3* ^34^, *foxj1a* ^35^, with digoxygenin RNA labelling kit (Roche DIG RNA Labeling Kit: 11175025910). In situ RNA hybridizations were performed as described ^36^.

### Immunohistochemistry and microscopy

Primary antibodies used for this study: acetylated tubulin (Sigma T7451: 1:500), Myosin 1d (Abcam ab70204: 1:400), atypical PKC (Santa Cruz sc-216: 1:200), Secondary antibodies were anti mouse-Alexa 594 (Invitrogen A11005: 1:500), rabbit-Alexa 488 (Invitrogen A11008: 1:500). Embryos were fixed in 4% PFA overnight and then dechorionated in 1×PBS. Embryos were permeabilized in 1×PBS with 0.5% Triton X-100 and treated with blocking solution containing 1×PBS, 10% sheep serum, 1%DMSO, 0.1% TritonX for an hour. Primary antibodies were diluted in fresh blocking buffer and incubated with embryos for at 4 °C overnight and then washed in 1xPBS with 0.5% Triton X-100 (3X washes, 30 minutes each) at room temperature.

### KV imaging and analysis

*Tg*(*dusp6:EGFP*)^*pt21*^ transgenic embryos at 2-3S were maintained at 28.5 °C and mounted on a transparent (MatTek: P35G-1.0-20-C) glass bottom culture dish in 1.5% low melting agarose. For CFTR expression analysis, KV was imaged in live *TgBAC*(*cftr-GFP*) embryos using a Perkin-Elmer UltraVIEW Vox spinning disk confocal microscope with 40x water immersion objective. Imaris (Bitplane) and Fiji (ImageJ NIH, Bethesda, MD) software was used to measure raw integrated density of the total KV area or the apical membrane region. The apical signal was then divided by the total signal to determine the apical intensity as a percentage of total fluorescence to control for differences in KV size or transgene expression levels. For 4D time-lapse imaging, we used Stimulation Emission Depletion (STED) imaging with Leica 93× 1.3 NA motorized correction collar objective. Z stacks with 200 nm step size imaged for 10 min with 8 sec intervals and were deconvolved with Huygens Professional version 17.04 (Scientific Volume Imaging, The Netherlands) in order to improve signal to noise ratio. All the measurements on speed and volume of vacuole was done manually from 4D STED images using Fiji. Vacuole volume measurements were derived from diameter, so only spherical structures were considered for measurement.

### Transmission Electron Microscopy

8S embryo samples were fixed overnight using cold 2.5% glutaraldehyde in 0.01 M PBS. Fixed samples were washed 3× in PBS then post-fixed in aqueous 1% OsO_4_, 1% K_3_Fe(CN)_6_ for 1 hour. Following 3 PBS washes, the pellet was dehydrated through a graded series of 30-100% ethanol, 100% propylene oxide then infiltrated in 1:1 mixture of propylene oxide: Polybed 812 epoxy resin (Polysciences, Warrington, PA) for 1 hr. After several changes of 100% resin over 24 hrs, pellet was embedded in molds for cross-sectioning embryos, cured at 37^°^C overnight, followed by additional hardening at 65^°^C for two more days. Ultrathin (70 nm) sections were collected on 200 mesh copper grids, stained using a Leica EM AC20 automatic grid staining machine with 2% aqueous uranyl acetate for 45 minutes, followed by Reynold’s lead citrate for 7 min. Sections were imaged using a JEOL JEM 1011 transmission electron microscope (Peabody, MA) at 80 kV fitted with a side mount AMT digital camera (Advanced Microscopy Techniques, Danvers, MA).

### Statistical analysis

Statistical significance using unpaired Students t-test and Mann Whitney’s test or one-way ANOVA and post hoc analysis using Turkey’s multiple range test by Graphpad Prism (Graph pad, La Jolla, CA, USA).

## ACKNOWLEDGEMENTS

We thank E. Korn (National Institute of Health, MD) for sharing *Acanthamoeba* sp. myosin-IC plasmid construct, S. Wolfe and N. Lawson (University of Massachusetts, MA) for their suggestions on *myo1d* targeting strategies, C. Stuckenholz and L. Davidson (Department of Bio Engineering) for sharing pCS2 CAAX and H2B: mCherry plasmid construct, and K. Alber (Center for Biological Imaging) at University of Pittsburgh, PA for help with tracking of vacuolar movement in time lapse images. This work was supported by NIH Grant 5R01GM104412 to MT and CWL. STED imaging instrument used for this work at CBI core facility was supported through NIH Grant S10 OD021540.

## AUTHOR CONTRIBUTIONS

CWL, HY, MS and MT initiated the project concept. MS and MT generated zebrafish *myo1d* mutants, and analyzed phenotypes. MS, MJC, JA and MT performed morpholino experiments. MS, TF, MSun and DS performed confocal, and TEM imaging and analysis. MS, MC, SW performed super-resolution 4D-STED imaging and analysis. MS analyzed all the data, prepared figures and drafted the manuscript. MT, JA, CWL edited the manuscript. All the authors have read the manuscript.

---

**Supplementary Figure 1.**
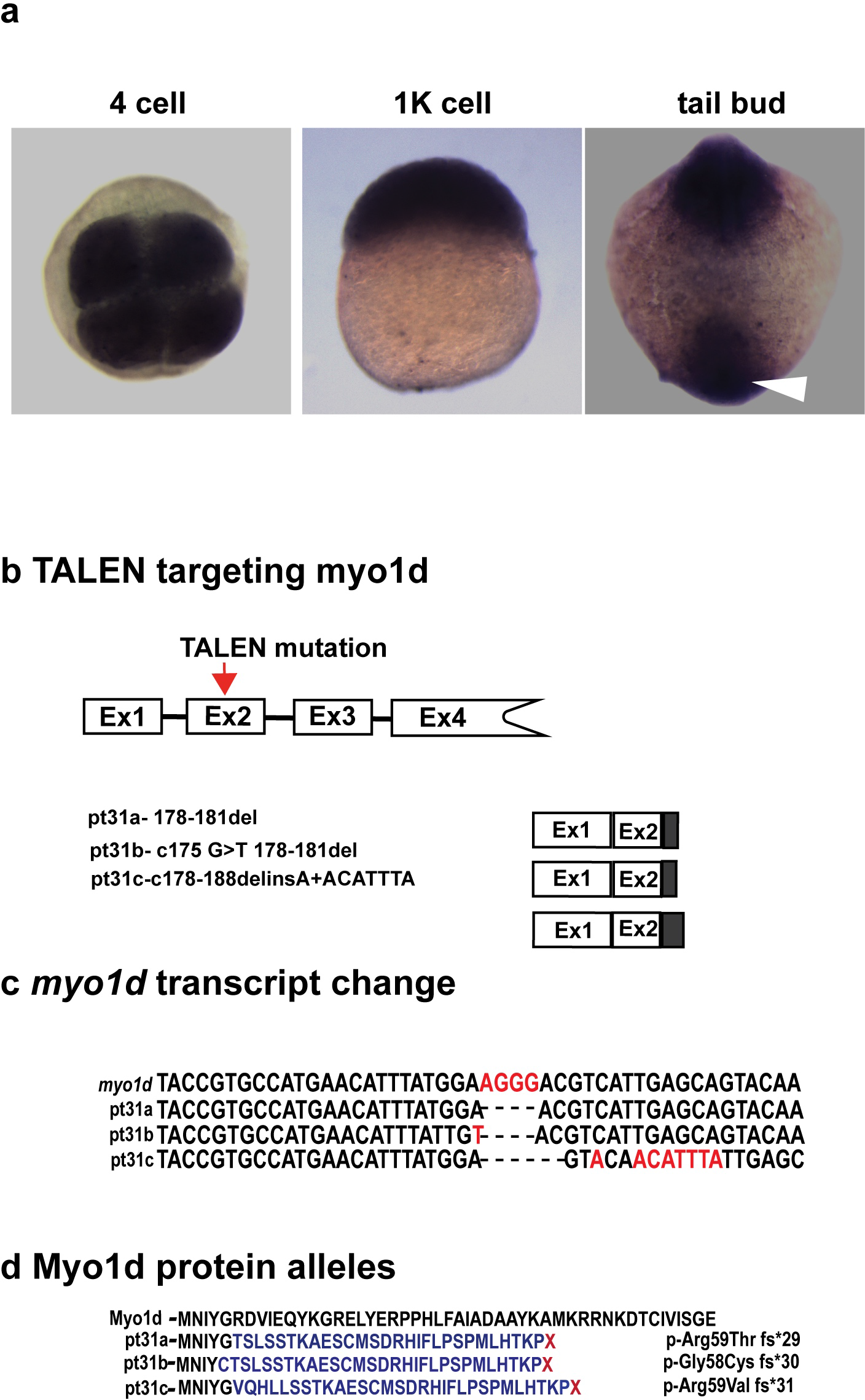
(a) mRNA expression of *myo1d* during early development. 4-cell, 1K cell and tailbud stages showed ubiquitous *myo1d* expression pattern. White arrowheads points to the region where KV is formed. (**b) *myo1d* targeting strategy in zebrafish**. A pair of goldy TALENs designed against *myo1d* exon 2. (**c**) Three F1 alleles identified, pt31a, b and c. (**d**) Predicted protein alleles show disruption of Myo1d by frame shift mutation and truncation.

**Supplementary Figure 2.**
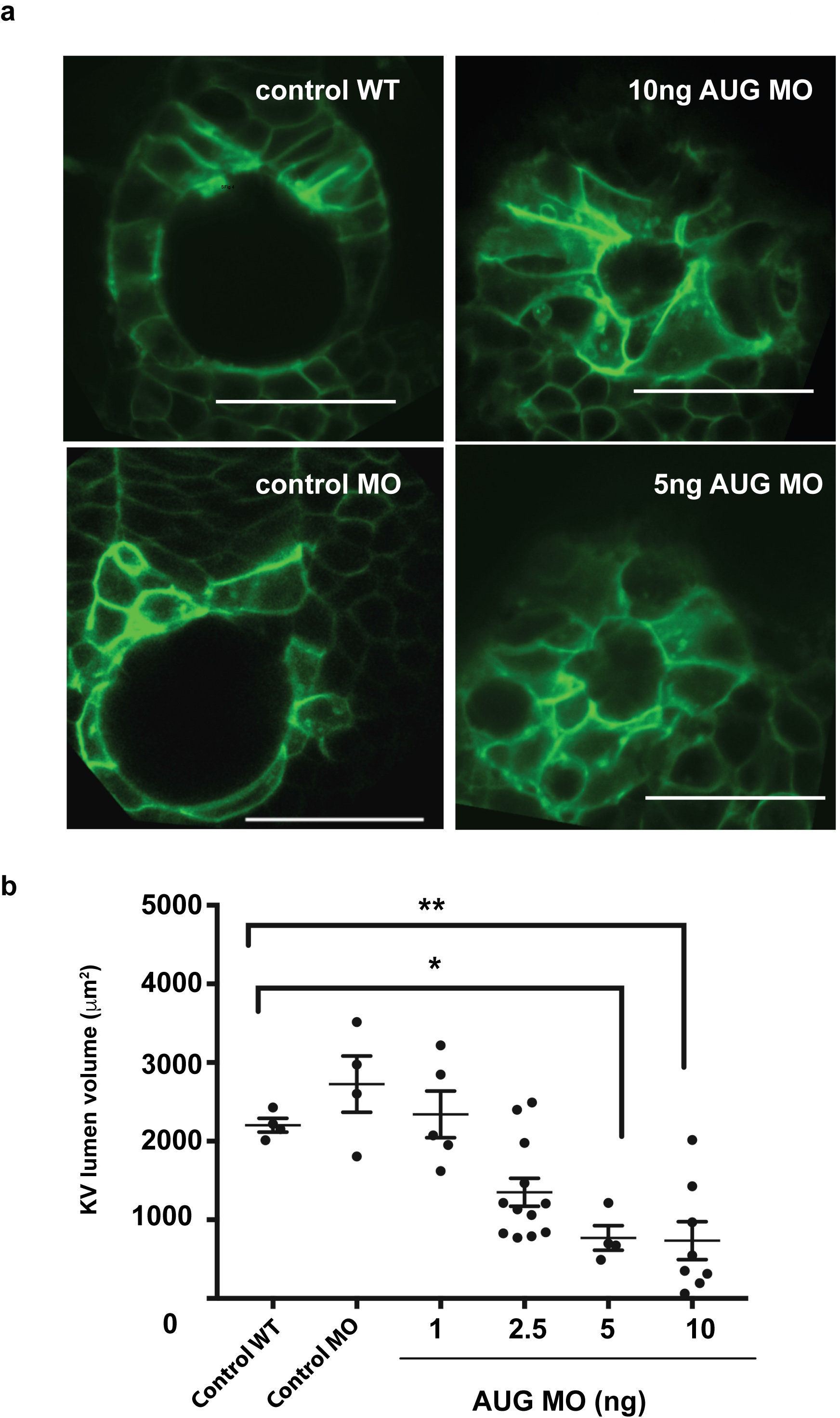
(a) Dose dependent reduction of KV lumen with *myo1d* AUG MO injection. Confocal Z plane images of KV showing lumen surface area with increasing doses of *myo1d* AUG MO injected into *Tg*(*dusp6:EGFP*)^*pt21*^ embryos. (**b**) Graph showing decreased lumen size with increasing doses of *myo1d* AUG MO injected embryos. wildtype (n=4), control MO (n=4), AUG MOs: (1ng, n=4), (2.5ng, n=12), (5ng, n=4), (10ng, n=8). scale bar-50jm. For **b**, one-way ANOVA and post hoc analysis with Turkey’s multiple range tests were used. *p<0.05, ** p<0.01 represent a statistical difference. **b**, Data as mean ± SEM. scale bar-50μm.

**Supplementary Figure 3.**
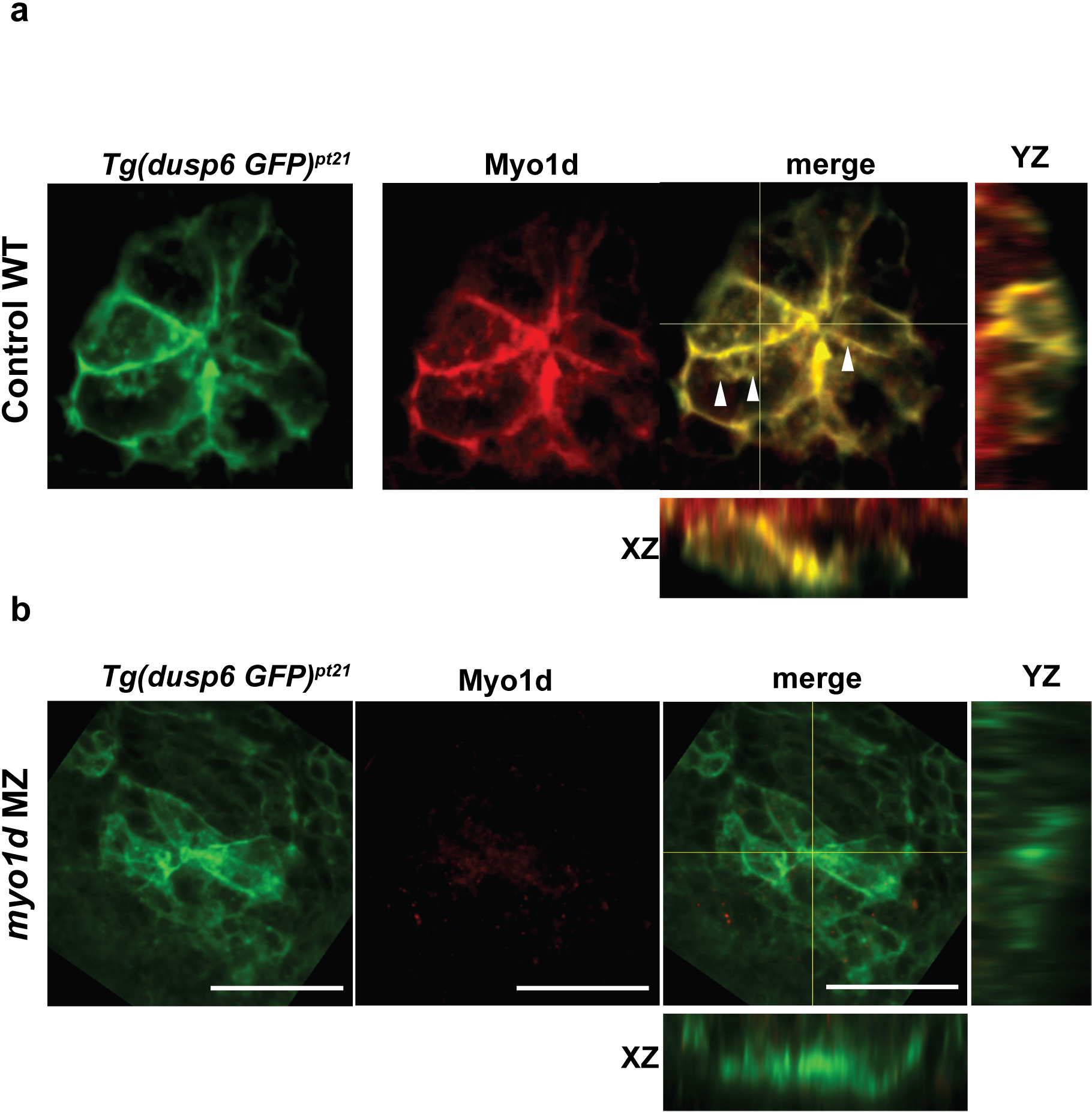
Myold expression in KV epithelial cells colocalized with vacuoles and was absent in *myo1d MZ* mutants. (**a**) *Tg*(*dusp6:EGFP*)^*pt21*^ embryos showing the expression of Myo1d (red) colocalized to the vacuoles in KV epithelial cells at 1 S (n=3). (**b**) Myo1d signal was absent in *myo1d MZ* (n=3). scale bar-50μm.

**Supplementary Figure 4.**
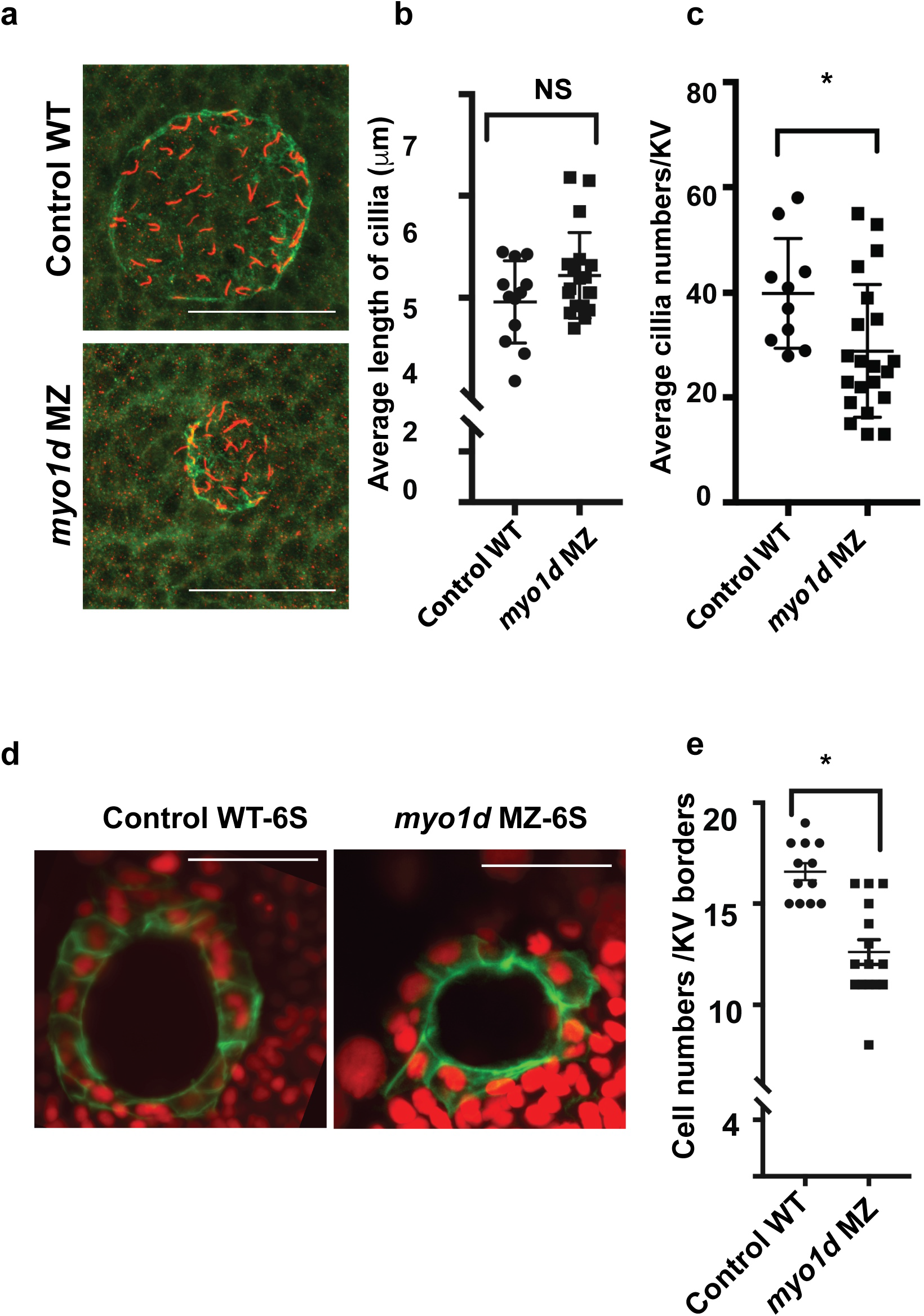
Ciliogenesis was unaffected in zebrafish KV. (**a**) Confocal 3D max projection showing KV size and cilia. 8S embryos stained for aPKC (green) and acetylated tubulin (red). (**b**) Average length of KV cilia was similar in wildtype (n=12) and *myo1d* MZ (n=19) embryos. (**c**) Average cilia numbers of KV were less *myo1d* MZ (n=19) than wildtype (n=12). (**d**) *Tg*(*dusp6:EGFP*)^*pt21*^ embryos in wildtype or *myo1d MZ* background injected with H2B mCherry mRNA showing KV epithelial cell morphology. (**e**) Cell numbers lining KV borders was decreased in *myo1d* MZ mutants (n=15) compared to wildtype (n=12). scale bar-50μm. Two sample comparisons in **b, c & e**, were by unpaired Students t-test and Mann Whitney test. Data as mean ± SEM. n represents number of KV analyzed. NS-Not significant, *p<0.05 represent a statistical difference.

**Supplementary Figure 5.**
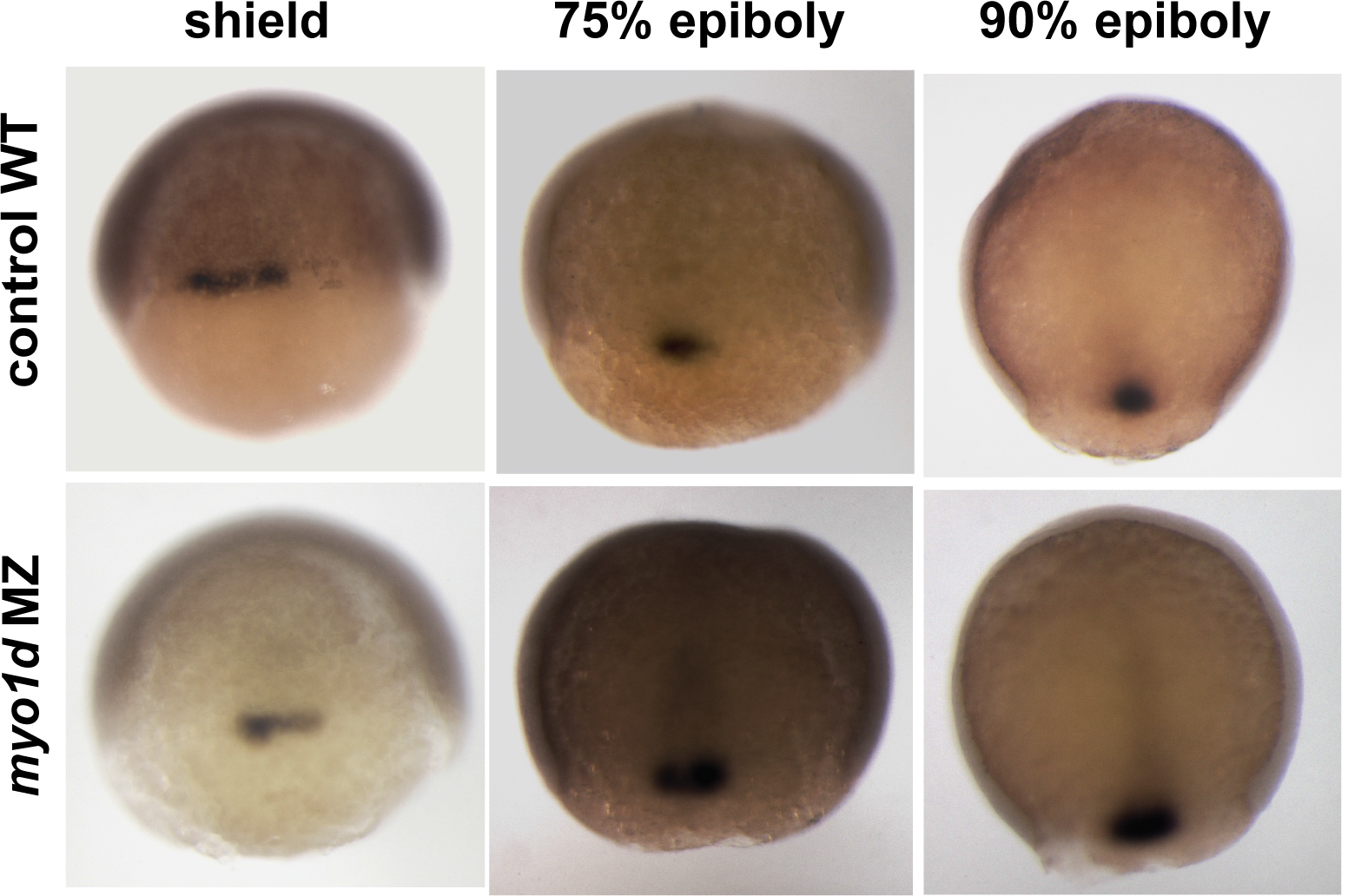
Dorsal Forerunner Cell (DFC) migration is not affected in *myo1d* MZ mutants. wildtype and *myo1d* MZ embryos were stained by *foxj1a* expression. Note only a single cluster of DFCs were detected at shield stage in both wildtype (22/36), and *myo1d* MZ, (29/36) embryos. This DFC cluster remained intact at 75% epiboly (wildtype (24/28), *myo1d* MZ (23/30)) and at 90% epiboly stages (wildtype (18/21), *myo1d* MZ (30/44)).

**Supplementary Figure 6.**
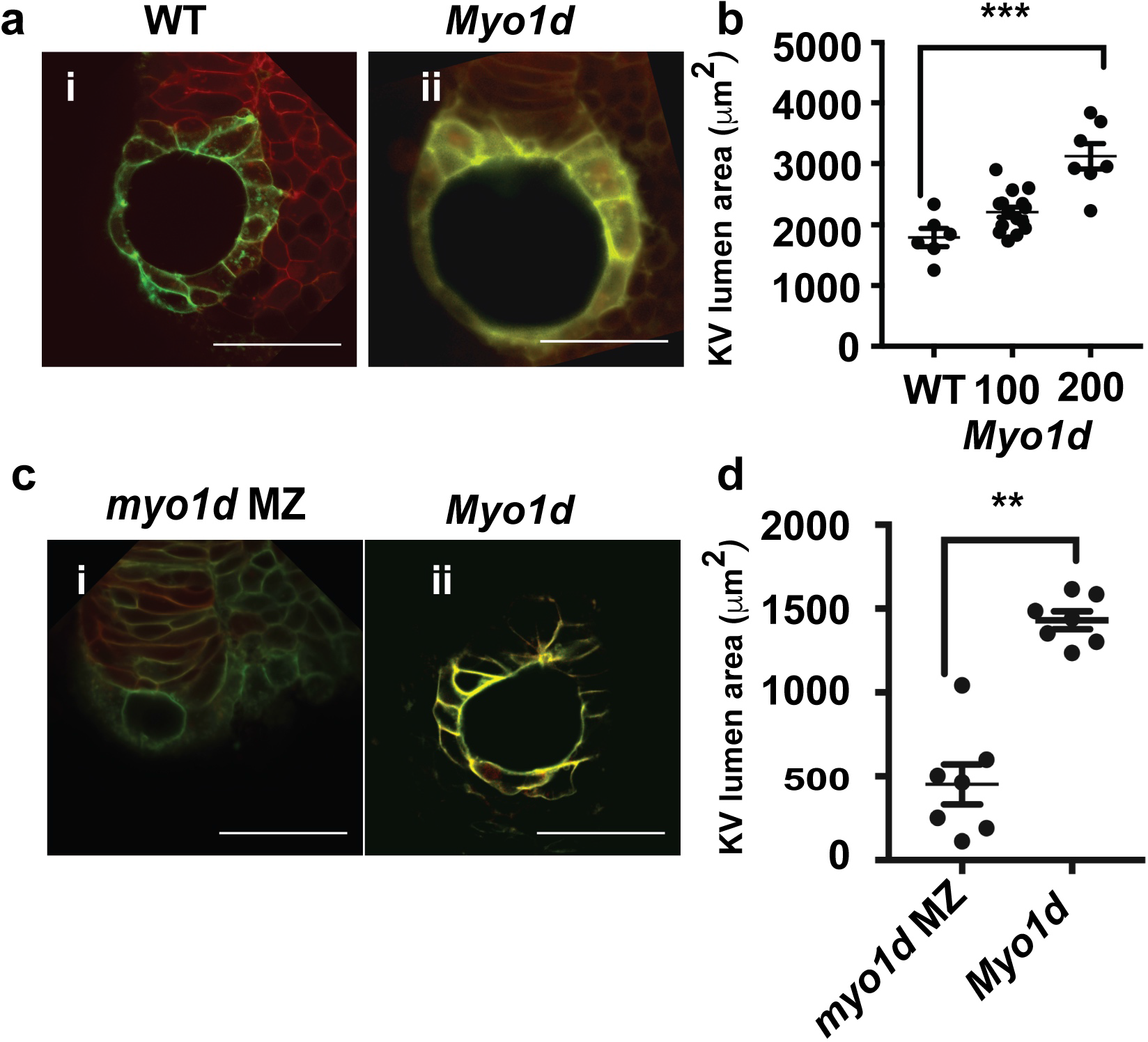
Expression of rat *Myold* mRNA rescued KV lumen defects in *myo1d MZ* mutants. **a**, Overexpression of rat MYO1D mRNA was sufficient to expand lumen area in wildtype embryos. **b**, Graph showing increase in lumen area due to overexpression of rat *Myo1d* (from 100pg (n=15) to 200 pg (n=7)) compared to wildtype uninjected KV (n=6). **c**, Overexpression of rat *Myo1d* mRNA rescued lumen formation defect in *myo1d* MZ mutants. **D**, Graph showing increase in lumen size after rat *Myo1d* overexpression (n=7) compared to uninjected *myo1d* MZ (n=7) embryos. For **b**, ANOVA and post hoc analysis with Turkey’s multiple range tests, whereas in **d**, unpaired Students t-test and Mann Whitney test were used. scale bar-50μm. Data as mean ± SEM. n-number of KV analyzed in the experiment. **p<0.001, ***p<0.0001 represent a statistical difference. 50 pg pCS2-CAAX mCherry DNA was used as injection control.

**Supplementary Figure 7.**
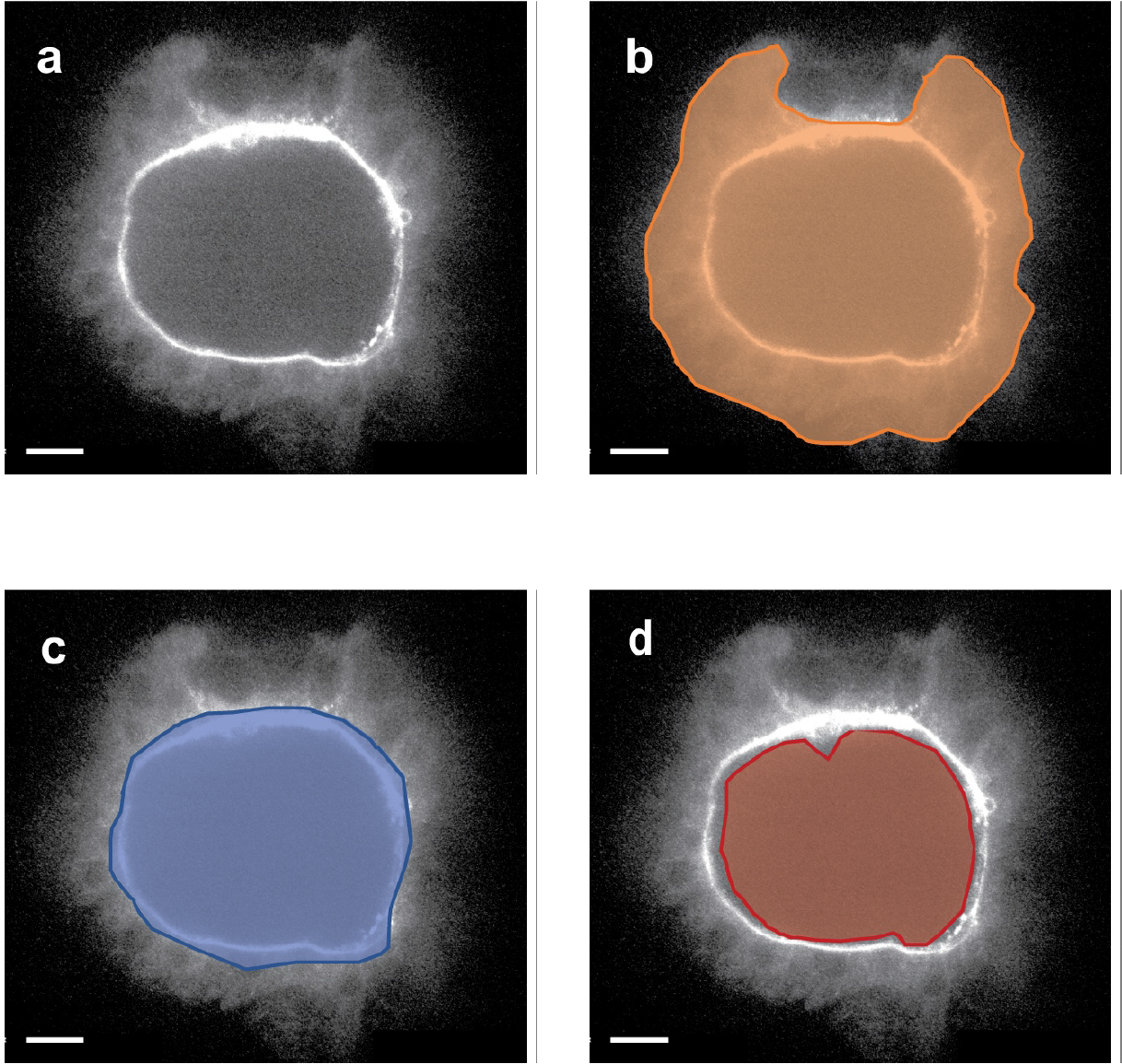
Quantification of CFTR apical expression. **a**, CFTR raw integrated density in 8S KV. **b**, Total KV area **c**, Apical area **d**, Lumen area.

**Supplementary Movie 1 Time lapse movie showing lumen expansion at 2-3 S in wildtype *Tg*(*dusp6:EGFP*)^*pt21*^ embryos**.

**Supplementary Movie 2 Time lapse movie showing lumen expansion at 2-3 S in *Tg*(*dusp6:EGFP*)^*pt21*^ embryos injected with *myo1d* MO.**

**Supplementary Movie 3 Confocal movie showing vacuolar transport across KV epithelial cells in wildtype embryos at 3S stage**. *Tg*(*dusp6:EGFP*)^*pt21*^ time lapse images were inverted for visualizing movement of vacuoles.

**Supplementary Movie 4 Confocal movie from another embryo showing vacuolar transport across KV epithelial cells in wildtype embryos at 3S stage.** *Tg*(*dusp6:EGFP*)^*pt21*^ time lapse images were inverted for visualizing movement of vacuoles.

**Supplementary Movie 5 Representative 4D STED Time lapse movie showing vacuolar transport across KV epithelial cells in wildtype embryos** (n=3). L-KV lumen

**Supplementary Movie 6 Representative 3D movie showing vacuolar lumen area and vacuole size across KV epithelial cells in *myo1dMZ* embryos** at comparable developmental stage (n=3). L-KV lumen

**Supplementary Movie 7 3D reconstructed movie showing vacuole predominance in posterior KV epithelial cells at 2-3 S stage in *Tg(dusp6:EGFP)*^*pt21*^ embryos.**

**Supplementary Movie 8 3D reconstructed movie showing vacuole predominance in posterior KV epithelial cells at 2-3 S stage in *myo1d* MZ embryos.**

**Supplementary Table 1.** Laterality score (expressed as percentage) in the myo1d AUG morpholino injection. Embryos fixed at 72hpf and probed using RNA in situ probes *(myl7, foxa3* and *spaw)*, scored for defective looping pattern.

|  | Control MO |  |  | *myo1d* MO |  |  |
| --- | --- | --- | --- | --- | --- | --- |
|  | left | straight+right | (n) | left | straight+right | (n) |
| *myl7* | 98 | 2 | 221 | 57 | 43 | 188 |
| *foxa3* | 87 | 13 | 120 | 72 | 28 | 276 |

|  | left | bilateral+right | (n) | left | bilateral+right | (n) |
| --- | --- | --- | --- | --- | --- | --- |
| *spaw* | 87 | 13 | 123 | 59 | 41 | 126 |

